# Illuminating links between cis-regulators and trans-acting variants in the human prefrontal cortex

**DOI:** 10.1101/2021.09.07.459322

**Authors:** Shuang Liu, Hyejung Won, Declan Clarke, Nana Matoba, Saniya Khullar, Yudi Mu, Daifeng Wang, Mark Gerstein

**Author notes:** Equal contributions (Shuang Liu, Hyejung Won, Declan Clarke).

## Abstract

Psychiatric disorders exact immense human and economic tolls in societies globally. Underlying many of these disorders is a complex repertoire of genomic variants that influence the expression of genes involved in pathways and processes in the brain. Identifying such variants and their associated brain functions is thus essential for understanding the molecular underpinnings of psychiatric disorders. Genome-wide association studies (GWASes) have provided many variants associated with these disorders; however, our knowledge of the precise biological mechanisms by which these contribute to disease remains limited. In connection with this, expression quantitative trait loci (eQTLs) have provided useful information linking variants to genes and functions. However, most eQTL studies on human brain have focused exclusively on cis-eQTLs. A complete understanding of disease etiology should also include trans-regulatory mechanisms. Thus, we conduct one of the first genome-wide surveys of trans-eQTLs in the dorsolateral prefrontal cortex (DLPFC) by leveraging the large datasets from the PsychENCODE consortium. We identified ∼80,000 trans-eQTLs. We found that a significant number of these overlap with cis-eQTLs, thereby implicating cis-mediators as key players in trans-acting regulation. We show, furthermore, that trans-regulatory mechanisms provide novel insights into psychiatric disease. Particularly, colocalization analysis between trans-eQTLs and schizophrenia (SCZ) GWAS loci identified 90 novel SCZ risk genes and 23 GWAS loci previously uncharacterized by cis-eQTLs. Moreover, these 90 genes tend to be more central in transcriptome-wide co-expression networks and more susceptible to rare variants than SCZ-risk genes associated by cis-variation.

## Introduction

Psychiatric diseases and neurological disorders afflict large portions of the global population and constitute a significant source of disability worldwide (*1*). Although genome-wide association studies (GWAS) have identified many genomic variants that are significantly associated with psychiatric and neurological disease risk, decades of research have led to only limited progress in our understanding of the precise mechanistic linkages between these variants and disorders of the brain.

Expression quantitative trait loci (eQTLs) are widely used to analyze the effects of variants on gene expression, i.e., an eQTL consist of an eSNP and an eGene where the eSNP affects the expression of the eGene. Previous studies have demonstrated that variants associated with human phenotypes frequently function as eQTLs [1], suggesting that eGene expression may play the role of so-called intermediary phenotypes. By definition, however, eQTLs are identified by statistical associations (i.e., correlation), whereas a fuller understanding of disease susceptibility and etiology entails better characterization of causal relationships. By investigating mediation effects in the context of eQTLs, we can more confidently establish these causal relationships.

It is now well established that cis-eSNPs may simultaneously be associated with distant genes [2,3]. A natural model for such phenomena may be one in which the expression of a trans-eGene (eGene affected by distant eSNPs) is mediated by cis-eGenes [2,4]. Trans-eQTLs that are mediated by cis-eGene expression provides more direct causal relationships between variants and trans-eGene expression. A simple but illustrative example of this phenomenon may involve an eSNP that lies within the promoter of a cis-eGene, wherein the cis-eGene is a transcription factor (TF). The distal target of this TF may then appear as a trans-eGene, the expression of which is strongly influenced by the variant affecting the expression of the regulator TF. Thus, the regulatory linkage of this example would be from the eSNP to the TF cis-eGene (mediator) to the trans-eGene. Elucidating the roles of mediators such as TFs, microRNAs, chromosomal remodeling proteins, and other regulatory factors provides immense value for understanding disease etiology in light of genomic variants. This immense value stems from the fact that a better characterization of mediators goes beyond statistical associations by providing a fuller picture of disease mechanisms via intermediary phenotypes, especially as they relate to trans-eGene expression [5,6]. Furthermore, uncovering instances of cis-mediation also enables investigators to elucidate and describe regulatory networks with greater confidence and thereby gain a more systems-level understanding of psychiatric diseases. Within this framework, for instance, cis-eGenes that regulate the expression of many trans-eGenes would function as so-called “cis-hub” genes [7,8], and these may form the basis of trans-eQTL hotspots within regulatory networks.

The Genotype-Tissue-Expression (GTEx) project has identified both cis- and trans-eQTLs in multiple human tissues [9]. Nonetheless, the sample sizes available for brain subregions were fairly limited in GTEx, thereby providing only a limited number of identified trans-eQTLs in brain tissues. In our previously published work on PsychENCODE data, we only calculated and reported cis-eQTLs.

In this paper, we combined the large-scale data resource generated by the PsychENCODE Consortium, CommonMind (CMC), and GTEx to identify trans-eQTLs with high confidence. We then carried out a careful set of analyses on the features of these trans-eQTLs and compared them with those of cis-eQTLs. Furthermore, our analyses enabled us to identify trans-eQTLs hotspots. By conducting both statistical mediation analysis as well as integrating data on inter-chromosomal contacts, we evaluated two potential mechanisms by which variants may influence trans-eGene expression. Upon integrating our results with schizophrenia (SCZ) GWAS, we demonstrated that trans-eQTLs might be pivotal for providing novel insights into disease mechanisms.

## Results

### Identification of trans-eQTLs in the human brain

Trans-eQTLs are especially difficult to identify in cohorts of limited sample sizes. To overcome this challenge, we worked with a large number of samples (N=1,387, which includes the PsychENCODE brain resource, CMC, and GTEx brain samples), thereby more readily enabling us to identify significant trans-eQTLs in a genome-wide fashion. We predicted that the resulting trans-eQTLs might reveal potential mechanisms of distal regulatory linkages across chromosomes. By adopting a standard approach for trans-eQTL identification that is similar to that of GTEx V8 [9], we tested associations between 12,245 highly-expressed genes and autosomal variants on a genome-wide scale (see **Methods** for processing and filtering criteria). We used the same covariates as those used for identifying cis-eQTLs in our previously published work [10]. Genes with poor mappability and variants located in repetitive regions were removed. Furthermore, trans-eQTLs between pairs of genomic loci with evidence of RNA-seq read cross-mapping were filtered out to minimize false positives [11]. We calculated FDR values from the LD-pruned list of trans-eQTLs to detect those that are significant. At an FDR threshold of 0.25, we detected 77,156 trans-eSNPs from ∼5.3M total SNPs tested in locations ! 5 Mb from the gene Transcription Start Site (TSS), comprising 17,899 independent SNPs after linkage-disequilibrium (LD) pruning. We identified 7,656 trans-eQTLs involving 582 eGenes at an FDR threshold of 0.05 (**Fig. 1A-B**). In summary, relative to previously published studies, we identified substantially more trans-eQTLs and trans-eGenes in the human brain by leveraging integrated data resources. We used the trans-eQTL list with FDR<0.25 for the analyses discussed in this study (Supplemental File 1). In addition to this primary list of trans-eQTLs, we also make available several lists of trans-eQTLs at varying FDR thresholds (**Fig. 1B**).

**Figure 1:**
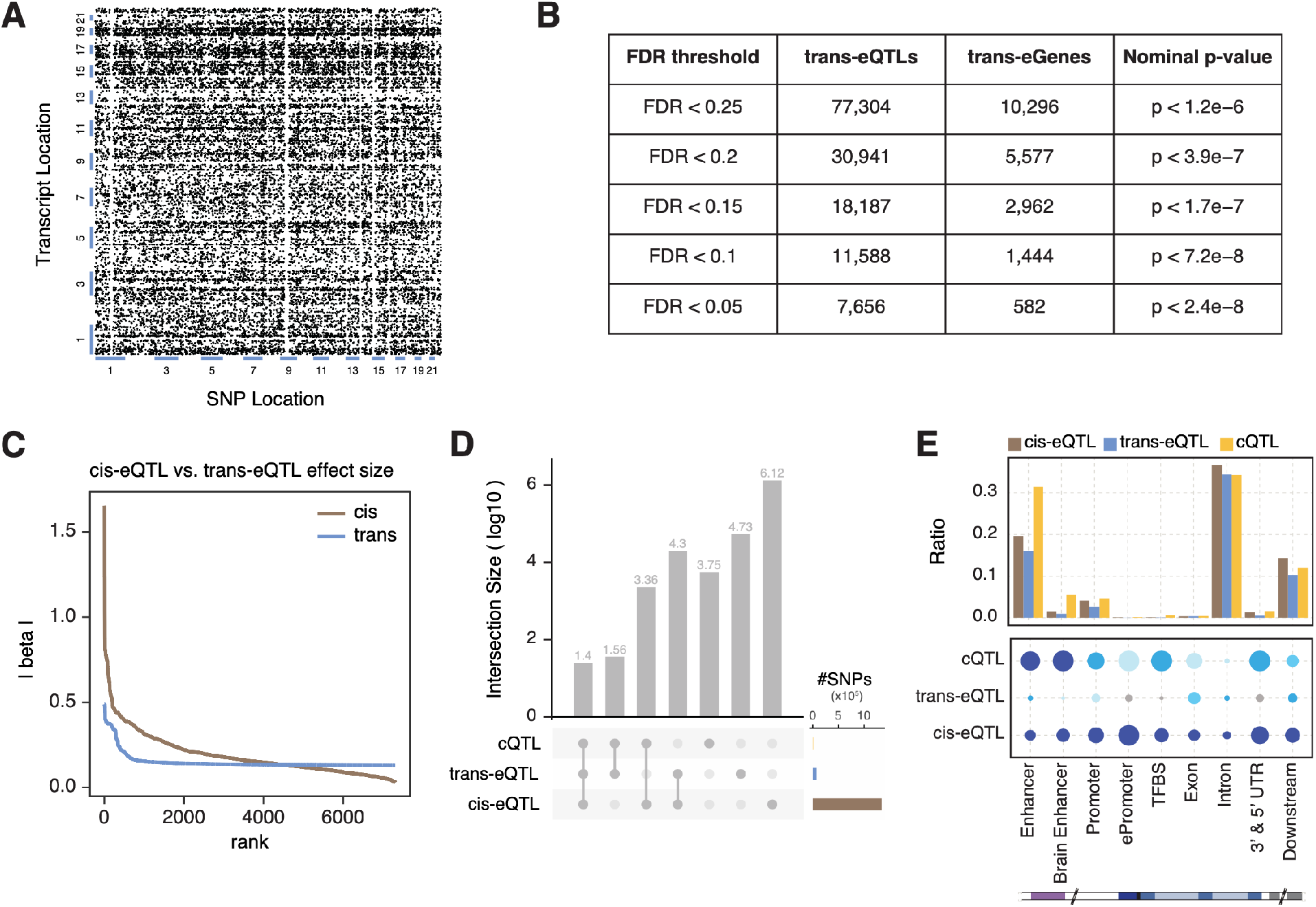
Characterization of trans-eQTLs. **A**. Genetic map for trans-eQTLs. **B**. Frequencies of trans-eGenes and trans-eSNPs at varying FDR thresholds. **C**. Comparisons of effect sizes between cis-eQTLs and trans-eQTLs. **D**. Frequencies of SNPs that are shared among various combinations of the 3 QTL types (cis-eQTLs, trans-eQTLs, cQTLs). **E**. Enrichment statistics and variant proportions are associated with the frequencies with which cis-eQTLs, trans-eQTLs, and cQTLs lie within various types of genomic elements.

We next characterized genomic features of trans-eQTLs and compared them with other types of QTLs in order to investigate associations between genomic elements and QTLs (**Fig. 1C-E**). In agreement with previous findings [12], we found that the magnitude of effect size for cis-eQTLs is larger than that of trans-eQTLs (**Fig. 1C**). Trans-eQTLs overlapped more frequently with cis-eQTLs than with cQTLs, which is also largely expected. We found 20,175 eSNPs shared between cis- and trans-eQTLs, whereas 61 SNPs were shared between trans-eQTLs and cQTLs. As shown in **Fig. 1D**, 25 SNPs were found to be shared among all 3 QTL types (i.e., cis-eQTLs, trans-eQTLs, and cQTLs). With respect to genomic elements, we found that trans-eQTLs tend to exhibit lower enrichment in most elements relative to cis-eQTLs and cQTLs. Trans-eQTLs were found to be most enriched within exons (**Fig. 1E**). The pattern of variant proportion on different genomic regions for trans-eQTLs is similar to that for cis-eQTLs but distinct from that of cQTLs.

### Potential mechanisms of trans-eQTLs

One potential mechanism by which specific variants exert trans-regulatory effects is one wherein the variant influences a nearby trans-regulator, which in turn regulates distal genes [2]. In this scenario, trans-eQTLs may also act as cis-eQTLs for nearby genes that have regulatory impacts on trans-eGenes (**Fig. 2A**). Indeed, we found that 19.33% of LD-pruned trans-eQTLs display cis-eQTL signals (**Fig. 2B**). Because simple genomic coordinate-level overlaps between cis- and trans-eQTLs may detect spurious associations due to LD, we performed a colocalization assay to identify cis- and trans-eQTL pairs that harbor shared causal variants. In total, we detected 1,688 trans-eQTLs (48.79% of trans-eQTLs that overlap with cis-eQTLs) that have shared causal variants with cis-eQTLs. We further interrogated the potential causal effects of cis-eQTLs on trans-eQTL associations via mediation analysis (**Fig. 2C**, Supplemental File 2). As part of this analysis, we found that 64.75% of trans-eQTLs that colocalize with cis-eQTLs can be explained by cis-mediators (p<0.05). Among these trans-eQTLs that can be explained by cis-mediators, there is roughly an even split of cis-mediators with positive and negative mediation effects, suggesting that trans-regulators can be either activators or repressors with roughly equal probability. As generally expected, we also observed that larger mediation coefficients tend to have greater statistical significance.

**Figure 2.**
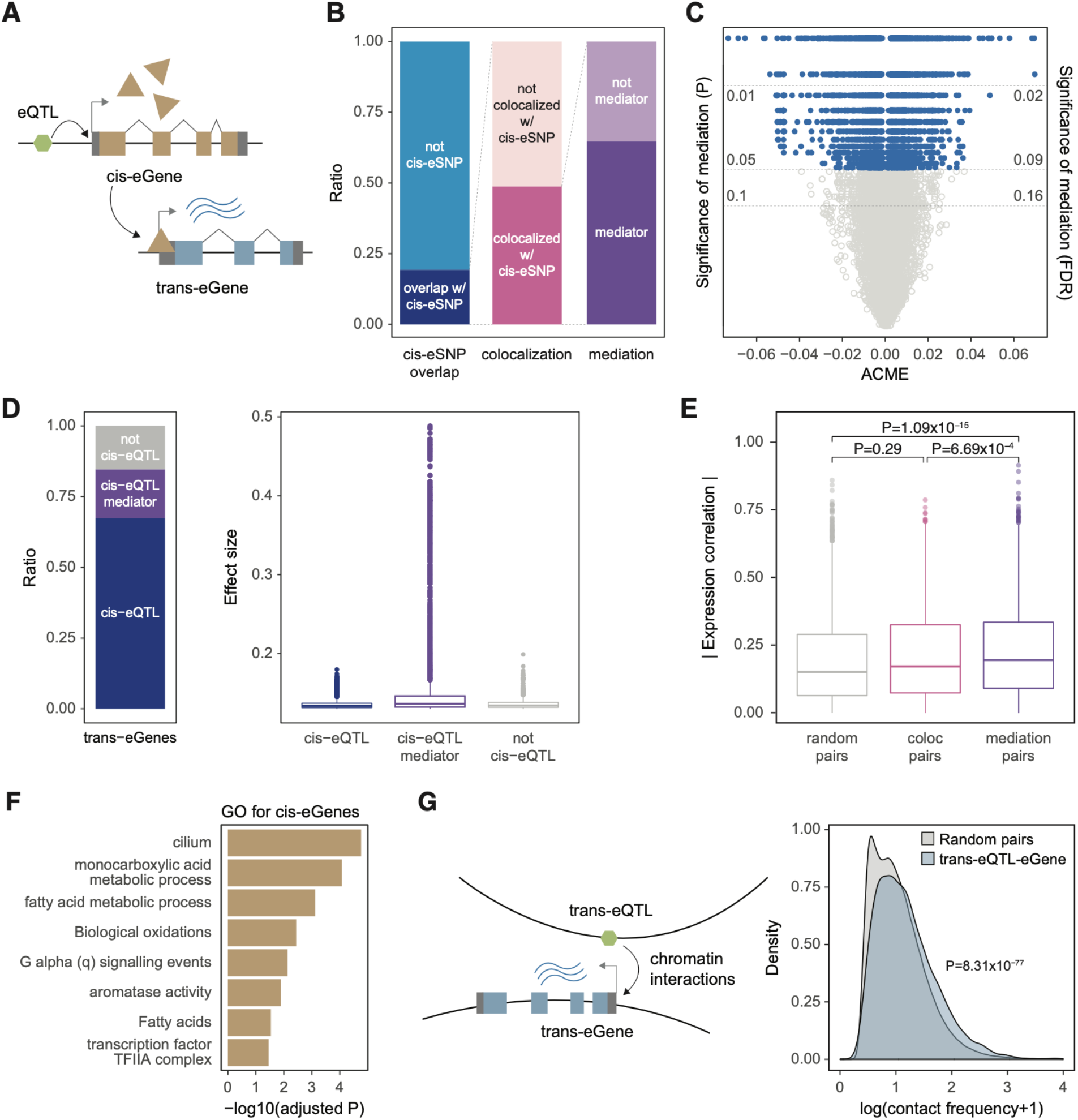
Cis-mediation and inter-chromosomal interactions explain trans-eQTL associations. **A**. A trans-eQTL may overlap with a cis-eQTL that regulates a proximal gene (cis-eGene), which may, in turn, regulate a distal gene (trans-eGene). **B**. Relative abundances of trans-eQTLs that overlap with cis-eQTLs (left), colocalize with cis-eQTLs (middle), and exhibit mediation effects (right). **C**. Roughly 65% of trans-eQTLs colocalized with cis-eQTLs exhibit mediation effects. The number of trans-eQTLs with positive mediation effects is almost the same as those that are negative. **D**. Roughly 77% of trans-eGenes are also cis-eGenes, and ∼20% of these have cis-mediators. Trans-eGenes with cis-mediators have larger absolute effect sizes. **E**. cis- and trans-eGene pairs with evidence of cis-mediation display significant co-expression compared to both random pairs and those pairs that exhibit colocalization but not mediation. **F**. Biological pathways associated with cis-eGenes that mediate trans-eQTL associations are enriched for metabolic processes and transcriptional regulation. **G**. trans-eQTL and eGene pairs tend to have higher inter-chromosomal interaction frequencies than do random pairs. ACME: for Average Causal Mediation Effects.

We found that 77% of trans-eGenes are also cis-eGenes, indicating that gene expression is regulated by both cis- and trans-acting variants. Roughly 20% of trans-eGenes had cis-mediators by cis-eGenes (**Fig. 2D**). Interestingly, the effect size of trans-eGenes with cis-mediators was larger than those without cis-mediators (p<2.2e-16, **Fig. 2D**).

By definition, because cis-mediation implies that variants influence trans-eGene expression via cis-eGenes, we hypothesize that cis- and trans-eGene pairs with evidence of cis-mediation are co-regulated. We evaluated this hypothesis by first grouping cis- and trans-eGene pairs into those that exhibit evidence of cis-mediation (which we term mediation pairs) and those with evidence of colocalization but mediation (which we term colocalization pairs). When comparing these groups, we found that mediation pairs showed greater expression correlation than colocalization pairs or expression-level matched random pairs (**Fig. 2E**; see Methods), thereby providing additional evidence for cis-mediation.

We next investigated the properties of cis-eGenes that were found to mediate trans-regulatory effects. In addition to being enriched for transcriptional regulators (e.g., members of TF complexes), cis-eGenes were also enriched for other biological processes (e.g., metabolic processes, **Fig. 2F**). These results indicate that variant effects on distal gene regulation are not solely dependent on TF activity. Instead, variants that are associated with metabolism may exert broad system-level effects on cellular function, which can then lead to changes in distal gene expression.

Another potential explanation for trans-eQTLs may lie in inter-chromosomal interactions. Previous studies have shown that inter-chromosomal interactions can bring multiple genes from different chromosomes into close physical proximity, thereby more easily enabling these genes to be co-regulated [13]. We, therefore, hypothesize that trans-eSNPs may regulate trans-eGenes located in different chromosomes via inter-chromosomal interactions. Indeed, we observed that trans-eSNP and eGene pairs display increased chromatin contact frequency compared with random inter-chromosomal contacts (**Fig. 2G**). Hence, in addition to cis-mediation, trans-eQTL associations can partly be driven by features of chromosomal conformation.

### Trans-eQTLs hotspots

Trans regulators exert broad impacts on the gene regulatory landscapes by affecting many downstream targets. Given the trans-regulatory properties of trans-eQTLs, some trans-eSNPs may affect multiple genes, thereby forming “trans-eQTL hotspots.” We defined such trans-eQTL hotspots as trans-eSNPs that affect three or more genes (**Fig. 3A**). In total, 382 trans-eQTL hotspots were detected. Because trans-eGenes for a given trans-eQTL hotspot are regulated by the same SNP, we expected that they might generally be co-regulated. Indeed, we found that trans-eGenes grouped by hotspots are significantly more co-regulated compared to expression-level matched random controls (**Fig. 3B**).

**Figure 3.**
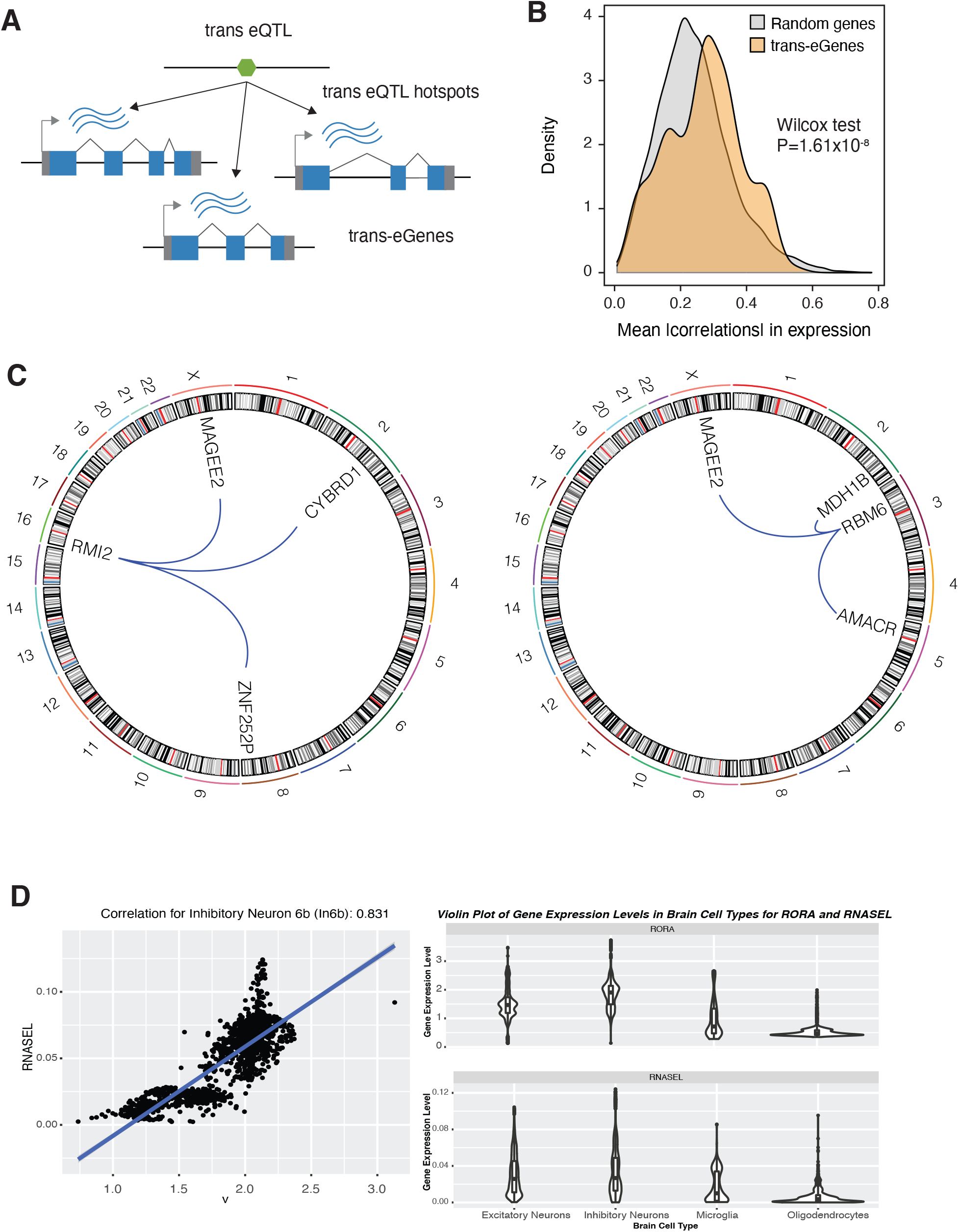
Trans-eQTL hotspots. **A**. Trans-eQTL hotspots represent trans-eSNPs that have associations with at least three trans-eGenes. **B**. Trans-eGenes associated with the same trans-eSNPs is co-regulated. **C**. Examples of trans-eQTL hotspots. **D**. Cell-type-specific TF-TG relationships detected by our mediator-trans-cis-QTL network.

One of the trans-eQTL hotspots consisted of three trans-eGenes *(MAGEE2, CYBRD1, ZNF252)*, which are distributed across the genome and are regulated by a trans-eSNP (chr16:11394372; **Fig. 3C**). Notably, this trans-eSNP was a cis-eQTL for *RMI2*, a gene associated with genome instability and Bloom syndrome [14]. In another example, the cis-eSNP for *RBM6* (chr3:50257020), an RNA binding protein, was associated with three trans-eGenes *(MAGEE2, MDH1B, AMACR*, **Fig. 3C**). Collectively, these results suggest that multiple biological processes (such as genome instability and RNA processing) may exert broad impacts on gene regulation via trans-regulatory mechanisms.

### Cell-type gene regulatory effects of trans-eQTLs and mediators

Because trans-eQTLs were defined from the brain homogenate and lack cell-type specificity, we evaluated cell-type-specific trans-regulatory effects from our mediators to trans-genes. For instance, our mediation analysis indicated that RORA (a nuclear receptor TF) is a mediator for the trans-eGene *RNASEL*, which encodes mammalian endoribonuclease. Based on single-cell multi-omics data [15], we found that RORA regulates the gene *RNASEL* specifically in neuronal cell types [15]. As shown in **Fig. 3D**, cellular expression levels of *RORA* and *RNASEL* show high Pearson correlation coefficients in several neuronal cell types, especially in inhibitory types, such as In6b (r = 0.831), In8 (r = 0.790), Ex9 (r = 0.746), In6a (r = 0.742), compared to glial cell types, e.g., microglia (r = 0.6436) and oligodendrocyte (r = 0.492).

Both genes are found to be involved in brain diseases. For example, the overactivation of RNASEL may be harmful in neurodevelopmental and inflammatory genetic diseases such as the Aicardi-Goutières syndrome [16]. RNASEL expression may result from signals from activation of NMDA receptors in cortical neurons by glutamate, and it may lead to the degradation of RNA molecules; degradation of mitochondrial RNA by RNASEL may contribute to neuronal death overall [17].

Decreased levels of nuclear receptor TF RORA have been found in the prefrontal cortex and cerebellar neurons of individuals with Autism Spectrum Disorder (ASD) [18]. Retinoic Acid signaling pathways are some of the neuronal circuits that are disrupted in ASD individuals [19]. In fact, the decreased expression of RORA impacts the regulation of its target genes in ASD individuals (several that are ASD-relevant genes) and tends to be associated with the pathobiology of ASD, such as decreases in neuronal differentiation and survival, poorer synaptic transmission, and neuroplasticity, worse cognition and spatial learning, memory impairment, and disrupted development of the cortex and cerebellum [18]. Furthermore, it has been found that RORA is involved in the differentiation of Purkinje cells, development of the cerebellum region, protection of neurons against oxidative stress, circadian clock rhythm regulation, and suppression of inflammatory processes [18].

### Trans-eQTLs identify novel disease mechanisms

It has been proposed that trans-eQTLs explain 60-90% of the heritability of gene expression [12,20]. However, functional annotation of GWAS variants largely relies on the use of cis-eQTLs, which may miss key biological underpinnings of human traits and disease. Trans-eQTLs may provide novel insights into the biological mechanisms underlying psychiatric illnesses. Motivated by these ideas, we performed colocalization analysis [21] between SCZ GWAS [22] and trans-eQTLs to unveil previously uncharacterized SCZ-associated biological pathways driven by trans-regulatory mechanisms (**Methods**). These results were then compared with the colocalization results between SCZ GWAS and cis-eQTLs. We found that some loci only colocalized with cis- or trans-eQTLs but some colocalized with both.

In total, we found that trans-eQTLs could explain 55 out of 142 SCZ-associated genome-wide significant (GWS) loci (**Fig. 4A**). In contrast, cis-eQTLs explained 78 GWS loci (**Fig. 4A**). Thirty-two GWS loci colocalized with both cis- and trans-eQTLs, suggesting that a subset of SCZ loci may exert their effects via multiple regulatory mechanisms. Furthermore, 23 GWS loci colocalized only with trans-eQTLs but not with cis-eQTLs, suggesting that trans-eQTLs may provide regulatory mechanisms for previously unexplained loci.

**Figure 4.**
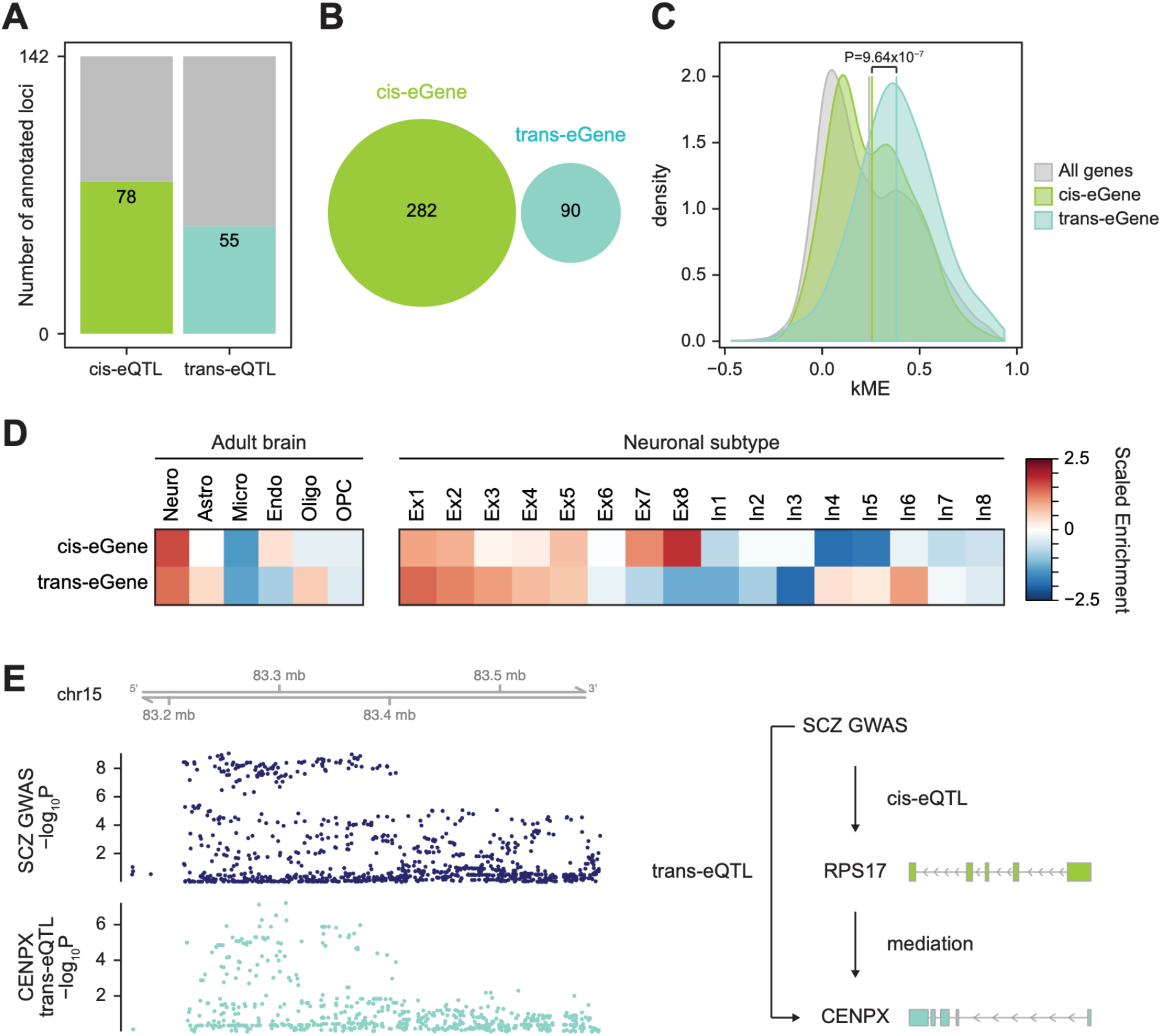
Trans-eQTLs identify novel SCZ risk genes and biological pathways. **A**. The number of SCZ GWS loci explained by cis- and trans-eQTLs. **B**. SCZ cis-eGenes do not overlap with SCZ trans-eGenes. **C**. Trans-eGenes that colocalize with SCZ GWAS exhibit greater network centrality. **D**. Cellular expression profiles of SCZ-associated cis- and trans-eGenes. **E**. An SCZ GWS locus colocalizes with trans-eQTLs for *CENPX* (left). This GWS locus is a cis-eQTL for *RPS17*, a cis-mediator of *CENPX* (right).

Colocalization analysis resulted in 90 and 282 SCZ-associated trans- and cis-eGenes; we refer to these as SCZ-(trans/cis)-eGenes, respectively (**Fig. 4B**). As expected, none of the SCZ-cis- and SCZ-trans-eGenes overlapped, demonstrating that trans-eQTLs can pinpoint distinct SCZ-associated genes and biological pathways. In particular, SCZ trans-eGenes were found to be enriched for JUN kinase activity (FDR=0.0056), a signaling pathway involved with neuronal apoptosis, neurite outgrowth, and dendrite arborization [23].

Notably, two of the 90 SCZ trans-eGenes *(AKAP11, SETD1A)* also harbor SCZ-associated rare variants *(de novo* loss-of-function (LoF) variation) [24] (Fisher’s exact test, P=0.0096, OR=14.62, 95% CI=1.67-59.06). This contrasts with SCZ cis-eGenes, none of 282 which overlapped with the genes that harbor SCZ-associated rare *de novo* LoF variants. This result corroborates the omnigenic hypothesis [12]. According to this hypothesis, genes can be divided into core genes that directly affect the disease-related biological processes and peripheral genes that regulate core genes. Core genes are likely to be affected by rare variants with large effect sizes, suggesting that these two SCZ trans-eGenes are likely core genes. Also, the hypothesis suggests that core genes are also likely to be targeted by trans-regulatory mechanisms, which is also consistent that these two genes are trans-eGenes of SCZ rare variants and mediated by peripheral genes.

To provide additional support for the omnigenic hypothesis, we explored the network properties of SCZ trans-eGenes. Core genes are thought to be enriched for network hubs [25]. Therefore, we measured module membership (kME; a measure of network centrality) of SCZ-eGenes in brain co-expression networks constructed from the same samples [26]. SCZ trans-eGenes showed significantly higher kME values than did cis-eGenes or brain-expressed genes in general (**Fig. 4C**, Two-Sample Wilcoxon test: cis vs. trans, P=9.64×10^−7^; all vs. trans, P=2.87×10^−9^; all vs. cis, P=0.099). These results demonstrate that SCZ trans-eGenes are characterized by greater network connectivity.

A significant portion of SCZ-associated trans- and cis-eGenes were found to be differentially regulated in brain tissue of SCZ-affected individuals [26] (Fisher’s exact test: trans, P=0.0094, OR=1.87, 95% CI=1.14-2.98; cis, P=0.035, OR=1.38, 95% CI=1.01-1.85). However, they were enriched in different SCZ-associated co-expression networks. SCZ-trans-eGenes showed selective enrichment in the SCZ-associated gene module 7 (geneM7, beta=0.0033, FDR=0.011), a neuronal module involved with synaptic vesicle formation, and which exhibits elevated expression signatures in SCZ (Fisher’s exact test: P=8.02×10^−5^, FDR=0.0014, OR=5.52, 95% CI=2.42-11.10). In contrast, SCZ cis-eGenes were only nominally enriched in the gene module 8 (geneM8, beta=-0.0030, FDR=0.017), a neuronal module downregulated in SCZ (Fisher’s exact test: P=0.018, FDR=0.53, OR=2.31, 95% CI=1.13-4.24). Therefore, SCZ-associated trans- and cis-eGenes may account for different expression signatures of SCZ.

Cell-type expression profiles of SCZ-eGenes illustrated potential cell types that contribute to distinct expression features (**Fig. 4D**). Both SCZ-associated trans- and cis-eGenes were highly expressed in neurons, which is consistent with previous findings that neurons are the primary cell type underlying SCZ etiology [27–29]. However, SCZ cis-eGenes exhibited relatively higher expression in lower layer neurons (Ex7-8), while SCZ trans-eGenes showed upper-to-lower layer gradient expression. Furthermore, while SCZ cis-eGenes were relatively depleted in inhibitory neurons, SCZ trans-eGenes were highly expressed in parvalbumin-expressing GABAergic interneurons (In6). In fact, recent studies have found that glutamatergic and dopaminergic dysfunction is associated with key symptoms of SCZ [30], and this may result from defects in neurotransmission by GABAergic interneurons. Notably, these neurons are responsible for inhibiting control of cortical and subcortical circuits and are dysfunctional in bipolar disorder as well as SCZ [31].

One example of a SCZ-trans-eGene is *CENPX*, which is associated with kinetochore assembly and DNA damage repair [32]. SCZ GWAS colocalized with trans-eQTLs for *CENPX*, which was mediated by cis-eQTLs for *RPS17*, a gene that encodes a ribosomal protein (**Fig. 4E**). This result suggests that an SCZ GWAS SNP may affect a nearby gene *RPS17*, which in turn regulates *CENPX*, a gene located in a different chromosome. Together, these results demonstrate how the integration of GWAS variants, cis-eQTLs, and trans-eQTLs can enhance our mechanistic understanding of disease etiology.

## Discussion

In this work, we report one of the first systematic searches and detailed studies of genome-wide trans-eQTLs in the dorsolateral prefrontal cortex (DLPFC). By leveraging the extensive resource built by the PsychENCODE consortium [10], we detected 77,304 trans-eQTLs (at an FDR threshold of 0.25) and 382 trans-eQTL hotspots. We reasoned that trans-eQTLs might provide new avenues for investigating trans-regulatory mechanisms, thereby providing critical insights for understanding the regulatory landscape in psychiatric disease and human health.

Coupled with increased multiple testing burdens, smaller effect sizes of trans-eQTLs make their identification considerably more challenging than the identification of cis-eQTLs. Indeed, we found that trans-eQTLs exhibit smaller effect sizes compared with cis-eQTLs. Furthermore, trans-eQTLs were less enriched in regulatory elements compared with cis-eQTLs, suggesting that trans-regulatory architectures may differ from those associated with cis regulation.

Despite these differences, a significant portion of trans-eQTLs overlapped with cis-eQTLs, suggesting that trans-acting variants may often exert their effects via cis-mediators. We found that trans-eGenes with evidence of cis-mediators displayed larger effect sizes and co-expression signatures with cis-mediators. Intriguingly, cis-mediators were involved with fatty acid metabolism, which contributes to ∼20% of the energy source of the brain [33]. This result indicates that trans-regulatory mechanisms may involve various biological processes that influence fundamental cellular function and psychiatric disease progression.

We also found that, in addition to cis-mediation, trans-eQTLs may also result from inter-chromosomal interactions. Understanding mechanisms that underlie trans-eQTL associations will provide important insights into trans-regulatory networks.

We further deconvolved trans-regulatory networks into specific cell-type-specific trans-regulatory networks and investigated the potential effects of the mediators we identified on trans-genes. These analyses uncovered a strong link between *RORA* regulation of *RNASEL*, both of which are associated with the immune system response and naive T cell states, and which also harbor polymorphisms associated with elevated cancer risk and mortality. Additional diseases beyond psychiatric disorders have been found to relate to *RORA* and *RNASEL*. For example, both genes have been shown to be associated with prostate cancer. In affected men, *RORA* is typically inactivated [34], and *RNASEL* has been considered to be a candidate for the hereditary prostate cancer gene *(HPC1)* [35]. In fact, there have been some mutations in the tumor suppressor gene *RNASEL* that lead to Ribonuclease L dysfunction, inflammation, infection, and increased risk of prostate cancer, suggesting links between innate immunity and tumor suppression [36]. These findings can help provide more insights into age-related changes, the roles of CFSs and genomic instability, the roles of the adaptive immune response (especially of T cells), and the potential roles of the central nervous system in the onset and progression of one of the world’s leading cancer types [37].

Given the differences between cis- and trans-eQTLs, we hypothesized that trans-eQTLs might provide novel insights into the biology of psychiatric disorders. Motivated by this notion, we related trans-eQTLs to SCZ GWAS variants to decipher trans-regulatory mechanisms that may contribute to SCZ etiology. Trans-eQTLs identified novel SCZ risk genes, a subset of which explained previously uncharacterized SCZ GWS loci. In total, trans-eQTLs explained 55 out of 142 SCZ GWS loci.

SCZ trans-eGenes differed from SCZ cis-eGenes in multiple respects. For example, SCZ- trans-eGenes included *SETD1A* and *AKAP11*, high-confidence SCZ risk genes that harbor rare *de novo* LoF variation [24]. In contrast, SCZ cis-eGenes did not overlap with any of the rare variation-targeted SCZ risk genes. *SETD1A* encodes a histone methyltransferase. Mice that carry an LoF mutation in *SETD1A* showed cognitive deficits, abnormal neuronal morphology, and transcriptional alterations, further implicating its role in SCZ etiology [38]. Moreover, we observed enhanced network centrality of SCZ trans-eGenes compared with SCZ cis-eGenes. This is consistent with our recent finding that hub genes were not enriched for schizophrenia heritability when cis-regulatory mechanisms were used to map genetic risk factors to genes [39]. These characteristics of SCZ trans-eGenes (e.g., overlap with rare variation and network centrality) correspond to the definition of the core genes from the omnigenic hypothesis [12] that are likley trans-regulated by rare variants, suggesting that trans-eQTLs may play crucial roles in understanding core biological principles of SCZ.

Cellular expression profiles further substantiated distinct biological processes represented by SCZ-associated trans- and cis-eGenes. While SCZ cis-eGenes were enriched in lower layer neurons, SCZ trans-eGenes were enriched in upper-layer neurons, suggesting that cis- and trans-regulatory mechanisms may influence distinct cortical circuitry. Moreover, SCZ trans-eGenes were enriched in parvalbumin-expressing interneurons, whose genetic and transcriptional association with SCZ were reported previously [26,40,41].

Other studies such as [42] have mapped eQTLs in individual cell types, and these have demonstrated the importance of individual cell subtypes in analyzing the effects of cell type-specific eQTL in other contexts, such as in human fibroblasts. In the example case of fibroblasts, this cell-type approach in capturing gene regulation has been demonstrated to be highly effective in that such regulations have not been captured by traditional bulk RNA-seq approaches. That study also performed induced pluripotent stem cell (IPSC) reprogramming, which may provide a future avenue of study. Another study [43] analyzed cell-type-specific eQTL models and performed genome-wide cis ct-eQTL analyses in the blood and brain, and identified the importance of myeloid cells in AD risk. The authors of that study identified several blood and brain AD biomarkers implicating microglia and the immune system in AD progression.

In conclusion, the transcriptional architecture of the human brain is orchestrated by both cis- and trans-regulatory variants, and trans-eQTLs can provide insights into disease biology that has been previously unexplored.

## Methods and Materials

### QTL analysis

We used the standard pipelines from ENCODE, GTEx, and other large consortia to uniformly process the raw sequencing data from PsychENCODE [44] (including RNA-seq and genotype data), as well as to identify functional genomic elements, such as brain enhancers, expressed genes, and eQTLs. We also processed other data types, such as Hi-C and single-cell data. We followed the GTEx pipeline for identifying all trans-eQTLs. We did this to ensure maximal compatibility between our results and our previously published cis-eQTL results and also to optimally enable comparisons between our results and those published previously. We found that lowly expressed genes that were not detected in a subset of samples can be falsely associated with SNPs due to the statistical fluctuation introduced by inverse quantile normalization. Therefore, genes that were not detected in any of 1,387 samples were removed from the analysis, leaving 12,278 genes. We used the QTLtools software package for trans-eQTL identification. Following the normalization scheme used by GTEx, the gene expression matrix was first normalized using quantile normalization, followed by inverse quantile normalization to map to a standard normal distribution (and to remove outliers). 50 PEER factors, genotype PCs, gender, and respective study were used as covariates in our calculations to identify cis-eQTLs. (Given our much larger sample size, we used considerably more PEER factors than GTEx.) For trans-eQTLs, we calculated the associations between gene expression and variants greater than the 5Mb window of each gene’s TSS (both upstream and downstream). These calculations were performed using genotype and gene expression data from 1,387 individuals (associations between a total of 12k genes and 5,312,508 variants were tested for potential QTLs).

We performed multiple testing corrections on nominal P-values by limiting FDR values to less than 0.05, 0.1, 0.15, 0.2, and 0.25 and generated several different lists of trans-eQTLs. We identified ∼77k trans-eQTLs involving ∼10K eGenes with FDR<0.25 (Supplemental File 1). We used this trans-eQTL list for the following analysis.

### Enrichment of genomic elements

We annotated SNPs of cis-eQTL, trans-eQTL, cQTL, and all SNPs used for QTL calculations to find the overlap of SNPs with genomic elements using SnpEff. We then tested the enrichment of the QTL SNPs in different genomic elements using Fisher’s exact test by using all SNPs used for QTL calculation as background. We also calculated the ratio of the number of SNPs in each genomic element to the total number of input SNPs. We selected Promoter, UTR, Exon, Intron, Downstream, TFBS, Enhancer, Brain Enhancer, and ePromoter regions for enrichment and ratio calculations.

### Mediation analysis

Trans-eQTLs that survived a genome-wide threshold of FDR<0.25 were further pruned for LD (r2>0.6), resulting in 74,143 independent trans-eQTLs. We overlapped trans- and cis-eQTLs that are in LD with a given independent trans-eQTL and ran colocalization analysis using the default settings of coloc [21]. Trans- and cis-eQTL pairs that survived colocalization analysis with the posterior probability (H4 PP, the posterior probability that indicates cis- and trans-eQTLs are sharing causal variants) greater than 0.5 were selected for mediation analysis. We used the SNP dosage and gene expression for the trans-eGenes and cis-eGenes associated with the SNP as input for the mediation analysis. We used the mediation package in R for the mediation analysis [45].

### Characterization of trans-eQTLs that have cis-mediators

We identified the trans-eGenes overlapped with of cis-eGenes, trans-eGenes not overlapped with cis-eGenes and cis-eGenes, which are mediators for trans-eGenes. We then compared the effect sizes of eGenes in these three categories using Kolmogorov–Smirnov test.

We hypothesize that cis-eQTLs may exert their effects on trans-eGenes via cis-eGenes that act as mediators. In this scenario, trans- and cis-eGene pairs that survive mediation analysis (mediation pairs) would need to be co-regulated. We, therefore, leveraged individual-level normalized expression data from Wang et al. [10] and measured expression correlation between cis- and trans-eGenes across 1,813 individuals. Because cis-mediator may act as both activators and repressors, we calculated absolute Pearson correlation coefficients associated with cis- and trans-eGene expression. Expression correlation coefficients of mediation pairs (mediation FDR<0.05) were then compared against expression the correlations of random gene pairs (random pairs) and colocalized, but not mediated cis- and trans-eGene pairs (coloc pairs). To generate random pairs, pairs of genes were randomly selected after matching for the expression level with cis- and trans-eGene pairs. Coloc pairs were identified as cis- and trans-eGene pairs that are colocalized (H4 PP>0.5) but do not show a sign of mediation (mediation P>0.1).

We also surveyed the function of cis-mediators via gene ontology (GO) analysis using gProfiler [46]. We used all cis-eGenes as a background gene list. GO terms that survived multiple testing corrections with the g:SCS threshold less than 0.05 were selected.

### Interchromosomal interactions

SNPs may affect genes located in another chromosome via interchromosomal interactions. Therefore, we quantified interchromosomal interactions between trans-eSNPs and trans-eGenes (trans-eQTL association FDR<0.25) using normalized chromatin contact matrices of the adult DLPFC [10] at 100kb resolution. Contact frequencies between trans-eSNPs and trans-eGenes were compared against contact frequencies between randomly selected 100kb bins with matching chromosomes. Since interchromosomal contact frequencies are often zero, we only retained and compared non-zero interaction frequencies between trans-eQTLs and random pairs.

### Trans-eQTL hotspots

Trans-eSNPs that are associated with 3 or more trans-eGenes were classified as trans-eQTL hotspots. Among 74,143 independent trans-eQTLs, 382 variants were identified as trans-eQTL hotspots. With the hypothesis that trans-eGenes associated with a given hotspot share a common trans-regulator, we evaluated whether trans-eGenes associated with hotspots may be more co-regulated than expected by chance. To this end, we calculated the mean absolute Pearson correlation coefficient among trans-eGenes associated with a given trans-eQTL hotspot using the individual-level normalized expression data from Wang et al [10]. Mean absolute expression correlation coefficients were compared between trans-eQTL hotspots and randomly selected expression-matched gene sets.

### Schizophrenia (SCZ) GWAS vs. trans-eQTL colocalization

Because the number of trans-eQTLs is much smaller than cis-eQTLs, we relaxed the thresholds for trans-eQTLs for colocalization analyses, and retained only trans-eQTLs with p<1e-5. We overlapped trans-eQTLs (here, trans-eQTLs with p<1e-5 were retained) with SCZ GWS loci (defined on the basis of LD [r2>0.6] with the index SNP) using the intersect function of bedtools. We then performed colocalization analysis between SCZ GWAS and trans-eQTLs using the default setting of coloc [21]. Trans-eQTLs that colocalized with SCZ GWAS at a threshold of H4 PP>0.6 were selected to identify trans-eGenes associated with SCZ (hereafter referred to as SCZ trans-eGenes).

To draw direct comparisons with SCZ trans-eGenes, SCZ cis-eGenes were defined using the same colocalization posterior probability as that we had used for SCZ trans-eGenes (H4 PP>0.6). We filtered previously defined SCZ risk genes [10] using a threshold of H4 PP>0.6.

### Characterization of SCZ trans-eGenes

Genes with excess of rare *de novo* LoF variation in SCZ were obtained from the SCHEMA browser [47]. 32 genes that showed significant association with SCZ at an FDR<0.05 [24] were overlapped with SCZ trans-eGenes. We performed Fisher’s exact test between genes with SCZ-associated rare variation and SCZ trans-eGenes. Protein-coding genes were used as a background list in performing Fisher’s exact test.

To interrogate potential dysregulation of SCZ-trans/cis-eGenes in SCZ-affected individuals, they were intersected with genes that were differentially expressed in postmortem brain samples with SCZ (SCZ-DEGs) or co-expression modules associated with SCZ (SCZ-modules) [26]. SCZ-DEGs were selected based on FDR<0.05 regardless of whether the genes were upregulated or downregulated in SCZ. We ran Fisher’s exact test between SCZ-trans/cis-eGenes and SCZ-DEGs/SCZ-modules with the background list defined as genes whose expression was detected in Gandal et al [26]. Because Gandal et al [26] defined 34 co-expression modules (20 of which are associated with SCZ), P-values for over-representation analysis between SCZ-trans/cis-eGenes and SCZ-modules were corrected for multiple testing.

Network connectivity (kME values) was also obtained from Gandal et al [26]. Each gene was assigned to a co-expression module via weighted gene co-expression network analysis (WGCNA) [48]. We quantified the kME value of a given gene in a co-expression module to which the gene belonged. We then compared kME values of SCZ trans-eGenes with kME values of SCZ cis-eGenes or all genes.

Next, we interrogated cellular expression profiles of SCZ-trans/cis-eGenes using single-cell RNA-seq data [10], as described previously [27]. We scaled expression profiles of each cell and calculated the average expression of SCZ-trans/cis-eGenes in a given cell. This cell-level expression value of a given gene was then aggregated based on the cell types (neurons, astrocytes, microglia, endothelial cells, oligodendrocytes, and oligodendrocyte precursor cells) or neuronal subtypes (namely, excitatory, and inhibitory neuronal subtypes).

### Trans-regulatory network linking variants and regulatory elements to genes

To analyze how eQTLs affect gene regulatory networks, we combined trans-eQTLs with gene regulatory networks from four brain cell types: neurons (excitatory and inhibitory) and glial cells (microglia and oligodendrocytes). We mapped both cis-eQTLs and trans-eQTLs onto the gene regulatory network, which may provide information about how eQTLs break certain TF binding sites (TFBSs) and thereby result in changes to gene regulatory networks.

For the cis-network, we used all genes in eQTL pairs (SNPs and Genes) as target genes (TGs). We filtered network edges by overlapping SNPs with binding sites on enhancers or promoters. We then evaluated whether these SNPs break the corresponding TFs using motifbreakR, which works with position probability matrices to interrogate SNPs for their potential effects on TF binding. We filtered the edges, thereby leaving only those edges that have SNPs with their target genes. The final results provided us with a data set that contains each edge representing SNPs -> TFs -> enhancer/promoter -> TGs linkages. We can visualize the results by plotting a genomic region surrounding certain SNPs, as well as potentially disrupted motifs.

Generating the trans-network entailed a similar procedure. However, for trans-eQTL pairs, the SNPs may not be on the same chromosome as their target genes. When overlapping SNPs on binding sites, this introduces many problems. Therefore, we used mediator-eQTL data instead of trans-eQTL data when filtering out edges. After we carried out the same analysis as that detailed above for the cis-network, we mapped mediators back to trans-eQTLs (along with their target genes) to generate the results associated with the trans-network.

As a result, we built a mediator-trans-cis-QTL Gene Regulatory Network (GRN) using mediators (TFs) and trans-network (TGs) cis-network results; this regulatory network links SNPs to mediators to the trans-genes that are regulated by them, respectively. Next, we utilized two types of predicted cell-type GRNs from scGRNom [15] for the major brain cell types: excitatory neurons (Ex1, Ex2, Ex3e, Ex4, Ex5, Ex6a, Ex6b, Ex8, and Ex9), inhibitory neurons (In1a, In1b, In1c, In3, In4a, In4b, In6a, In6b, In7, and In8), microglia, and oligodendrocytes. The first predicted GRN type corresponds to cell-type open chromatin regions from scATAC-seq data. The second predicted GRN type corresponds to a filtered network with the top 10% TFs that have absolute coefficients for a given target gene regardless of whether the region is characterized by cell-type-specific open chromatin. We then overlaid these TF-TG edges in both versions of cell-type GRNs with those in our mediator-trans-cis-eQTL network to determine cell-type specific TF-TG relationships detected by our mediator-trans-cis-eQTL network.

## Supplementary information

Supplemental File 1 – trans-eQTLs with FDR<0.25 in DLPFC

Supplemental File 2 – mediation analysis results for trans-eQTLs

## Author contributions

D.W. and M.G. conceived and designed the study. S.L., H.W. and D.C. analyzed the data and presented the results. N.M., S.K. and Y.M. contributed to the data analysis. All authors wrote, read, and approved the manuscript.

## Acknowledgments

This work was supported by National Institutes of Health grants, R01AG067025 (D.W.), R21CA237955 (D.W.), R03NS123969 (D.W.), DP2MH122403 (H.W.), R21DA051921 (H.W.), R00MH113823 (H.W.), U01MH122509 (H.W.), U01MH116492 (M.G.).

